# Mass media promotion of a smartphone smoking cessation app: Modeled health and cost impacts

**DOI:** 10.1101/486316

**Authors:** Nhung Nghiem, William Leung, Christine Cleghorn, Tony Blakely, Nick Wilson

**Author notes:** Corresponding author: Prof Nick Wilson.

## Abstract

**Background:** Smartphones are increasingly available and some high quality apps are available for smoking cessation. However, the cost-effectiveness of promoting such apps has never been studied.

**Objective:** To estimate the health gain, inequality impacts and cost-utility from a five-year promotion campaign of a smoking cessation smartphone app compared to business-as-usual (no app use for quitting).

**Methods:** A well-established Markov macro-simulation model utilising a multi-state life-table was adapted to the intervention (lifetime horizon, 3% discount rate). The setting was the New Zealand (NZ) population (N = 4.4 million). The intervention effect size was from a multi-country randomised trial: relative risk for quitting at six months = 2.23 (95%CI: 1.08 to 4.77). Intervention costs were based on NZ mass media promotion data and the NZ cost of attracting a smoker to smoking cessation services ($NZ64 per person).

**Results:** The five-year intervention was estimated to generate 6760 QALYs (95%UI: 5420 to 8420) over the remaining lifetime of the population. For Māori (Indigenous population) there was 2.8 times the per capita age-standardized QALYs relative to non-Māori. The intervention was also estimated to be cost-saving to the health system (saving NZ$115, 95%UI: 72.5 million [m] to 171m; US$81.8m). The cost-saving aspect of the intervention was maintained in scenario and sensitivity analyses where the discount rate was doubled to 6%, the effect size halved, and the intervention run for just one year.

**Conclusions:** This study provides modelling-level evidence that mass-media promotion of a smartphone app for smoking cessation is likely to generate health gain, reduce ethnic inequalities in health and save health system costs. Nevertheless, there are other tobacco control which generate considerably larger health gains and cost-savings such as raising tobacco taxes.

## INTRODUCTION

Mobile phone-based interventions for smoking cessation have been shown to be effective, as per a Cochrane systematic review [1] and a meta-analysis [2]. Less research has been performed around the use of smoking cessation apps on smartphones. Nevertheless, a review of eight studies of such apps indicated favourable quit rates for app users in the range of 13% to 24% [3]. Others subsequent studies have also reported favourable results [4, 5], but the first full randomized controlled trials (RCT) were not published until 2018. One of these was a multicountry study (Australia, Singapore, UK and the US), that reported a quit rate at six months of 8.5% in the intervention group vs 3.8% in the control group (a relative risk [RR] = 2.23; 95%CI: 1.08 to 4.77; intention-to-treat analysis) [6]. A particular strength of this study was the comparison of a “state-of-the-art” decision-aid design versus support with passive information-only apps. Another RCT was conducted in Canada with 19 to 29 year-olds, and used a printed self-help guide for the control group [7]. It reported continuous abstinence at six months was not significantly different at 7.8% for the smartphone app versus 9.2% the self-help guide (odds ratio; OR 0.83, 95% CI 0.59-1.18). The third RCT was conducted in the US and involved mindfulness training via a smartphone app with experience sampling vs a control of experience sampling only [8]. It reported no group difference in smoking abstinence at six months (9.8% vs 12.1% in the two groups respectively, p = 0.51). But within the intervention group, the relationship between craving and cigarettes per day decreased as treatment completion increased (*p* = 0.04).

Promotion of smoking cessation using mass media campaigns (and more targeted advertising) has been found to be a cost-effective investment in tobacco control [9, 10]. In New Zealand (NZ), such mass media campaigns have also been reported to be cost-effective when promoting the national quitline service [11]. This suggested to us the possible value of promotion of smoking apps as an additional intervention for those smokers not using the quitline. Therefore we aimed in this modeling study to estimate the health gains and impact on health costs of this particular approach to tobacco control in New Zealand. This is a high-income country with a national Smokefree Goal for 2025 [12].

## METHODS

There seemed too much heterogeneity in the design of the three published RCTs to combine the results in a meta-analysis. So we just selected the RCT which we considered the most appropriate for the New Zealand population ie, the multi-country one) [6]. This RCT had a wider age range than the Canadian RCT and also the design of the control group was probably better than the Canadian trial (which used a printed self-help guide as opposed to another type of smartphone app in the multi-country trial). It would also be likely to have more general population appeal than the mindfulness app used in the RCT in the US.

Our modeled intervention consisted of promotion of the same particular app as used in the multicountry RCT [6], “Quit Advisor Plus” which is available for downloading free of charge from the Apple App Store. We assumed that for Android smartphone users, a similar quality and free smartphone app would be available, given our published survey of the quality of free smoking cessation apps available to New Zealand citizens on both Apple and Android platforms [13].

We assumed that these apps would be promoted on New Zealand health agency websites eg, the Ministry of Health (MoH) website. The website promotion was assumed to be at the level of $72,000 per annum, which is the amount spent on this purpose by the New Zealand Health Promotion Agency on its “Breakfast eaters campaign” (see Table S1 in Appendix). The apps would also be promoted with annual mass media promotion which was assumed to cost NZ$2,791,000 (the equivalent of the New Zealand quitline service’s marketing budget) [14].

We then used the average cost of attracting a Quitline caller to the New Zealand Quitline Service as a proxy for getting a person to download the smoking cessation app onto their smartphone (ie, NZ$64 per person [14]). This amount is fairly similar to the cost from a US study where mass media promotion was estimated to trigger app downloads at the advertising cost of ~NZ$70 per enrollee [15]. To estimate the number of smokers downloading the app, we divided the total expenditure on website promotion and mass media promotion by this $64 per person figure. This gave an estimate of 44,700 New Zealand smokers ($2,863,000/$64) who would be expected to download the app in response to the promotional activities. We included uncertainty around these estimates, along with age-variation in the download rate, albeit based on UK data for smoking cessation app downloads [16]. This gave download proportions by smoker age-group of: 11.3% for 20-34y; 9.0% for 35-54y; and 1.8% for 55+y (see Appendix for details).

The intervention effect size amongst those downloading the app was based on the only RCT with published results to date, as detailed in the *Introduction* (ie, relative risk [RR] for prolonged abstinence at six months = 2.23 (95%CI: 1.08 to 4.77) [6]. For simplicity, we assumed that only those who would have made an unaided cessation attempt download the app, ie, we assumed this was a different population from those who would tend to use the Quitline for quitting. Taking the 20-34-year-old smokers of European/other (non-Māori) ethnicity as an example, we estimated the annual net cessation rates for unaided quitting ie, at 3.1% (men) and 3.7% (women) (see Appendix, Figure S1, Tables S2 to S5). To this we applied the intervention effect size of RR = 2.23 amongst those estimated to be using the app in this age-group (11.3%), giving permanent quit rates of 7.1% (men) and 10.4% (women) for those app users in this demographic group (see Appendix for workings and results around quit rates for all the demographic groups eg, Māori ethnicity, 35-54y and 55+y age-groups). This heterogeneity was also included in the comparator arm of the model, where we also estimated the baseline unaided quit rates (ie, those not using the Quitline who are the target and comparator population for this evaluation).

In modeling health gain and net health system costs we used a well-established Markov macrosimulation model utilizing a multi-state life-table approach: the “BODE^3^ Tobacco Model” including probabilistic uncertainty about multiple input parameters [11, 17–21]. This model includes 16 tobacco-related diseases using national data by sex, age and ethnicity for the whole New Zealand population in 2011. It uses a health system perspective and estimates quality-adjusted life-years (QALYs) gained and net health system costs over the remainder of the population’s lifetime with both discounted at 3%. We also conducted sensitivity and scenario analyses around the discount rate, the effect size, impact for Māori (Indigenous population) and how many years the intervention was run for.

### Data availability

Supporting information with additional methods and results is attached. Data sharing with other researchers or official agencies of the precise data used in modeling is potentially possible subject to agreement with the government agencies making it available to the researchers (the Ministry of Health).

### Ethical approval

Approval for use of anonymized administrative data as part of the BODE^3^ Programme has been granted by the Health and Disability Ethics Committees (reference number H13/049).

## RESULTS

The five-year promotion of smoking cessation apps was estimated to generate 6760 QALYs (95%UI: 5420 to 8420) over the lifetime of the population (Table 1). For Māori there was 2.5 times the per capita gain relative to the non-Māori (at 3.14 vs 1.25 per 1000 population respectively); or a 2.78 times difference when age-standardized (Table 1 and Table S6). The intervention was also estimated to be cost-saving to the health system (saving NZ$115 million [m], 95%UI: 72.5m to 171m; US$81.8m (US$ for 2017). The overall cost-saving aspect of the intervention was maintained in all sensitivity and scenario analyses including where the effect size was halved (Table S7), the discount rate varied, including up to 6% (Tables S8 and S9), and the intervention run for just one year (Table S10). Other scenario analyses included running the intervention for 10 years and for 20 years (the latter yielding 19,600 QALYs gained and $418m in cost-savings, Tables S10 and S11). Key driver of uncertainty for both the health gain and cost-savings was the relative risk for net annual cessation rates comparing those who used the smartphone app to those who quit unaided (as per the tornado plots: Figures S2 to S5).

**Table 1.**
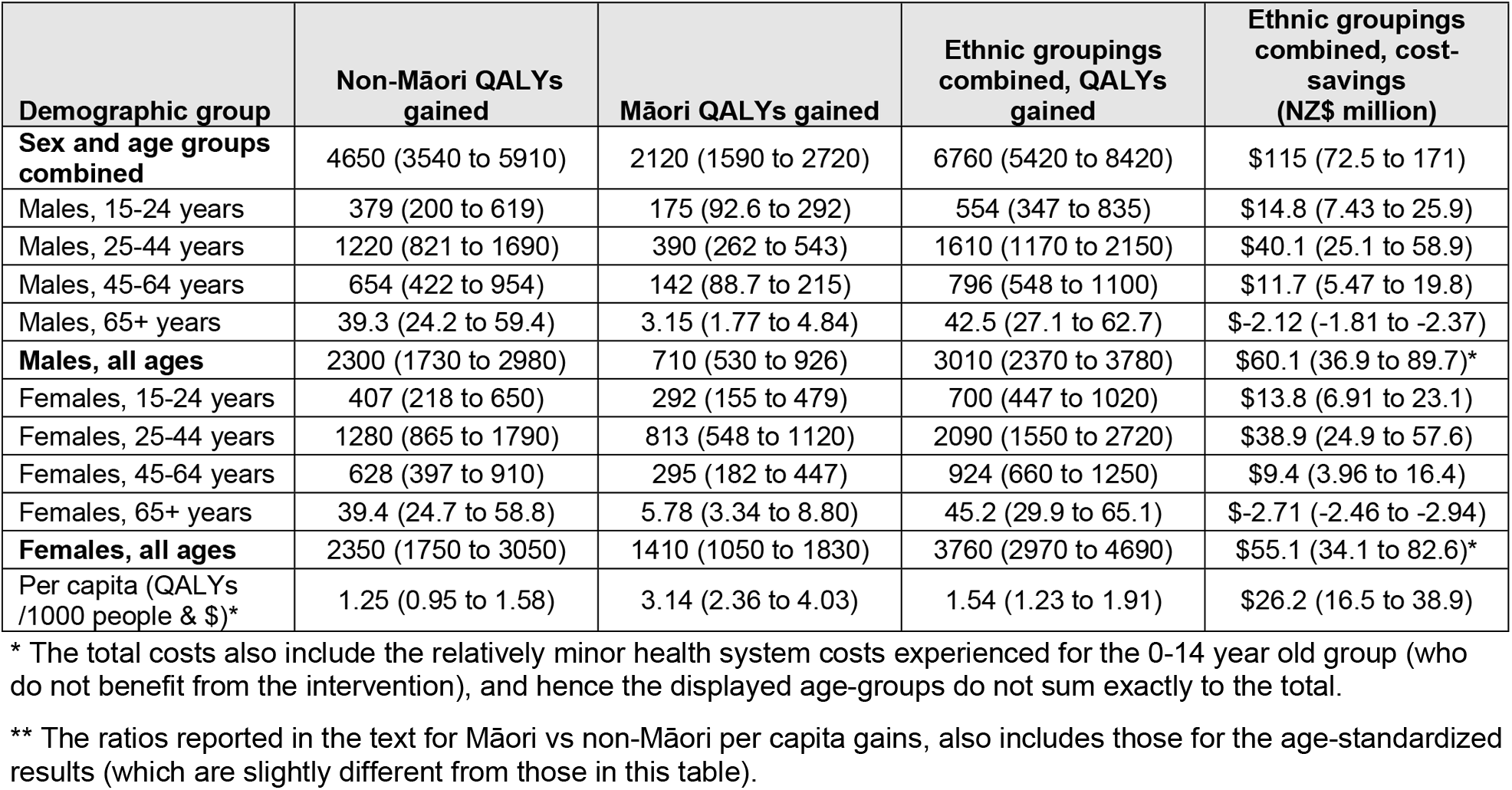
Health gain (QALYs) and health system cost-savings from the promotion of a smartphone app for smoking cessation (base case: effect size from a RCT, five-year intervention period, 3% discounting; 95% uncertainty interval shown in brackets)

## DISCUSSION

This modeling study suggests that the mass media promotion of a smartphone app and its use would be likely to generate overall health gains, favor greater per capita gains for the Indigenous population (Māori), and achieve cost-savings to the health sector. These findings are perhaps not surprising given the overall evidence for such apps being effective (see *Introduction*) and the evidence from a systematic review that mobile phone-based interventions [1] are effective. The cost finding is also consistent with a study showing that text messaging for smoking cessation is cost-saving for the health sector [22].

A strength of this modeling is that it is the first such study to consider the cost-utility of smoking cessation apps (not just text message interventions) and it uses a well-established tobacco control model containing detailed epidemiological and costing data. Nevertheless, there are various limitations with this modeling:

- The effect size was based on just a single RCT (with quitting measured at six months), albeit the RCT (out of the three published as of October 2018) to be considered the most relevant to the New Zealand population. However, we did a scenario analysis with half the effect size, which may better reflect the mixed outcome of these three trials.
- The baseline quit rate does not capture recent features of the tobacco control scene in New Zealand such as: the rise of e-cigarette use [23], the adoption of standardized tobacco packaging in New Zealand in 2018, and tobacco industry actions (eg, discount brands) that may undermine the ongoing annual tobacco tax increases used in New Zealand [24].
- No account was taken of potentially more efficient marketing strategies eg, via use of social media as per one study [25]. Similarly, no account was made of the potential synergies that could be achieved if app promotion was focused around the timing of World Smokefree Day activities or the annual rise in tobacco taxes in January of each year in New Zealand.

Given these issues, additional RCTs are desirable, especially those measuring quitting out to 12 months and actual costs of recruitment that might likely occur in population roll out. Furthermore, policy-makers need to compare app promotion with other potential tobacco control interventions eg, as per the published league table of tobacco control studies using the BODE^3^ Tobacco Model” in Nghiem et al [11] and online [26]. Some of these major interventions (eg, further tax increases, a tobacco-free generation and a sinking lid on supply) would be likely to generate much greater health gain (although they are also typically applied for greater than this five-year intervention, indeed lifelong), as well as accelerating progress to tobacco endgame goals.

## Competing interests

The authors declare no competing interests.

## Acknowledgements

The authors acknowledge colleagues who have helped with other aspects of building the tobacco control model used in this study: June Atkinson, Dr Linda Cobiac, Dr Giorgi Kvizhinadze.

## Funding

This work was supported by funding from the Ministry of Business, Innovation and Employment (MBIE), grant number: UOOX1406. Work on the original model was supported by a grant from the Health Research Council of New Zealand (grant 10/248).

## SUPPORTING INFORMATION

S1 Appendix

